# Dynamic redistribution of eIF4F controls cap-dependent translation initiation

**DOI:** 10.64898/2026.06.21.733607

**Authors:** Riley C. Gentry, Nicholas A. Ide, Victoria M. Comunale, Colin Echeverría Aitken, Colin D. Kinz-Thompson, Ruben L. Gonzalez

**Affiliations:** Department of Biological Sciences, Columbia University, New York, NY, USA; Laboratory of Molecular Biophysics, The Rockefeller University, New York, NY, USA; Department of Genetics, Harvard Medical School, Boston, MA, USA; Department of Chemistry, Columbia University, New York, NY, USA; Biochemistry Program, Vassar College, Poughkeepsie, NY, USA; Biology Department, Vassar College, Poughkeepsie, NY, USA; Department of Chemistry, Rutgers University-Newark, Newark, NJ, USA

## Abstract

Translation initiation requires messenger RNAs (mRNAs) to be recognized and loaded into ribosomes through a process catalyzed by the heterotrimeric eukaryotic initiation factor eIF4F. During this process, eIF4F engages the 7-methylguanosine cap at the 5’ end of the mRNA and promotes productive engagement with the ribosomal pre-initiation complex (PIC) to facilitate PIC loading onto the mRNA. Although eIF4F is central to translation initiation and its regulation, the molecular mechanism by which eIF4F stimulates PIC loading, and the mechanistic role of the essential ATP hydrolysis step catalyzed by eIF4F, have remained unresolved. Here, we use single-molecule fluorescence microscopy to directly visualize the dynamics of eIF4F during cap recognition and PIC engagement. We show that ATP binding, but not ATP hydrolysis, promotes productive assembly of eIF4F on mRNA and enables dynamic redistribution of eIF4F along the transcript. In contrast, ATP hydrolysis is specifically required for recycling of cap-stalled eIF4F during productive PIC engagement. Furthermore, we identify eIF3 and eIF4B as the minimal PIC-associated factors required to stimulate ATP-hydrolysis-dependent recycling of eIF4F during PIC loading. Together, our results support a model in which productive PIC engagement stimulates ATP-hydrolysis-dependent recycling of eIF4F, thereby coupling eIF4F recycling to PIC loading during translation initiation. This mechanism provides a framework for understanding how mRNA topology, RNA-binding proteins, and the availability of initiation factors can control translational efficiency.

## Introduction

Translation initiation constitutes a major regulatory stage of gene expression, determining when and which messenger RNAs (mRNAs) are translated into proteins. For more than 95% of eukaryotic mRNAs, translation initiation proceeds through the heterotrimeric eukaryotic initiation factor (eIF) 4F complex, composed of eIFs 4A, 4E, and 4G, which recognizes the terminal 5’ 7-methylguanosine cap (cap) present on nearly all cellular mRNAs^1–4^. Through a series of poorly understood steps, cap-bound eIF4F licenses the mRNA for loading into the mRNA-binding channel of the ribosomal 43S pre-initiation complex (PIC), which subsequently scans along the mRNA in the 3’ direction until it recognizes a start codon^1–4^. Collectively, these steps constitute the mRNA recruitment cascade. Consistent with its central role in translation initiation, eIF4F activity is tightly regulated; perturbations in eIF4F abundance are sufficient to drive widespread translational reprogramming, and numerous protein binding partners modulate eIF4F function through mechanisms that remain largely unresolved^1,5–7^.

Despite its fundamental role in gene expression and decades of investigation, the molecular mechanism by which eIF4F promotes translation initiation remains unresolved^1–4,8–23^. Using single-molecule fluorescence resonance energy transfer (smFRET), we recently defined the mechanism by which eIF4F recognizes the cap and primes mRNAs for translation^24^. These experiments demonstrated that eIF4F non-specifically samples the body of the mRNA and is actively displaced from cap-distal regions by its eIF4A subunit, enabling eIF4F to ‘hop’ along and dynamically redistribute throughout the mRNA until it becomes kinetically trapped at the cap (cap-stalled) through inhibition of eIF4A following cap recognition by eIF4E^24^. These observations led us to hypothesize that productive translation initiation would ultimately require release and recycling of the cap-stalled eIF4F complex, thereby enabling productive PIC engagement during loading onto the mRNA. They further implied that the essential ATP hydrolysis step catalyzed by eIF4F might function in one or more of these processes^3,20,22,23^. How ATP hydrolysis, eIF4F release, and productive PIC engagement are mechanistically coupled during translation initiation remains unknown.

Multiple competing models have been proposed to explain how eIF4F stimulates PIC loading and whether eIF4F remains associated with the PIC during subsequent stages of translation initiation^8,14,19–21^. Neither the timing nor the functional role of the essential ATP hydrolysis step catalyzed by eIF4F has been experimentally established^3,12,20,23^. Furthermore, although eIF4F is generally assumed to function as an RNA helicase that engages a complete 43S PIC, the specific PIC-associated factors that stimulate eIF4F activity, the mechanistic basis of that stimulation, and the composition of the PIC that is competent for productive engagement with eIF4F have not been experimentally determined^22,25–27^. The importance of resolving these questions is underscored by recent *in vivo* observations suggesting that a single eIF4F complex can drive multiple PIC-loading events, thereby directly modulating protein output from a given mRNA^28^. These uncertainties persist in large part because translation initiation substeps cannot be directly resolved *in vivo* and because most structural and biochemical approaches report on downstream mRNA-bound 48S PICs after eIF4F has already acted^8,9,19,22,25–27^.

To address these questions and define the molecular mechanism by which a cap-stalled eIF4F promotes translation initiation, we extended our recent *Saccharomyces cerevisiae* smFRET platform^24^ to directly monitor the dynamics of cap-stalled eIF4F during PIC loading onto the mRNA. Using fluorophore-labeled eIF4A and eIF4G together with ATPase-defective eIF4A mutants, we show that ATP binding regulates productive eIF4F assembly and identify the precise step at which ATP hydrolysis by eIF4F is required. Furthermore, by systematically varying PIC composition, we define the minimal set of eIFs required to couple eIF4F activity to productive PIC engagement. Together, these findings establish a new mechanistic model for cap-dependent translation initiation in which productive engagement between the PIC and cap-stalled eIF4F stimulates ATP-hydrolysis-dependent recycling of eIF4F, thereby mechanistically coupling eIF4F recycling to PIC loading and controlling translational output.

### ATP gates eIF4F complex assembly on mRNA

Since successful mRNA recruitment requires ATP hydrolysis by eIF4F, defining the timing and mechanistic role of nucleotide binding and hydrolysis is essential for understanding how eIF4F promotes translation. To define how ATP regulates eIF4F activity, we used the *S. cerevisiae*-based smFRET platform that we recently developed to define the mechanism of cap recognition by eIF4F^24^ (Figure 1a). This system monitors the position and dynamics of eIF4F along native mRNAs by measuring FRET between a Cy3 donor fluorophore positioned at a defined location on the *S. cerevisiae* rpl41a mRNA (Cy3-rpl41a) and a Cy5 acceptor fluorophore on the eIF4G subunit of eIF4F (Cy5-eIF4G). The assay is performed using wide-field total internal reflection fluorescence microscopy in a microfluidic flowcell, enabling simultaneous imaging of hundreds of individual Cy3-rpl41a mRNAs together with rapid exchange of solution conditions to inject additional proteins or eject unbound factors during both pre-steady-state and steady-state kinetic experiments. For these experiments (Figure 1a), Cy3-rpl41a mRNAs were tethered to the surface of the flowcell and interrogated with eIF4F complexes containing Cy5-eIF4G. Binding of eIF4F near the fluorophore-labeled position on the mRNA produced a high FRET efficiency (E_FRET_), whereas displacement or dissociation of eIF4F from such positions resulted in near-zero E_FRET_ (Figure 1a). Using this platform, we previously demonstrated that eIF4A functions as an ATP-dependent ‘strippase’ that removes eIF4F from uncapped mRNAs and from cap-distal regions of capped mRNAs, but not from the 5’ end of capped mRNAs^24^. Importantly, experiments using slowly hydrolyzing and non-hydrolyzable ATP analogs suggested that this stripping activity required ATP binding but not ATP hydrolysis^24^.

**Figure 1.**
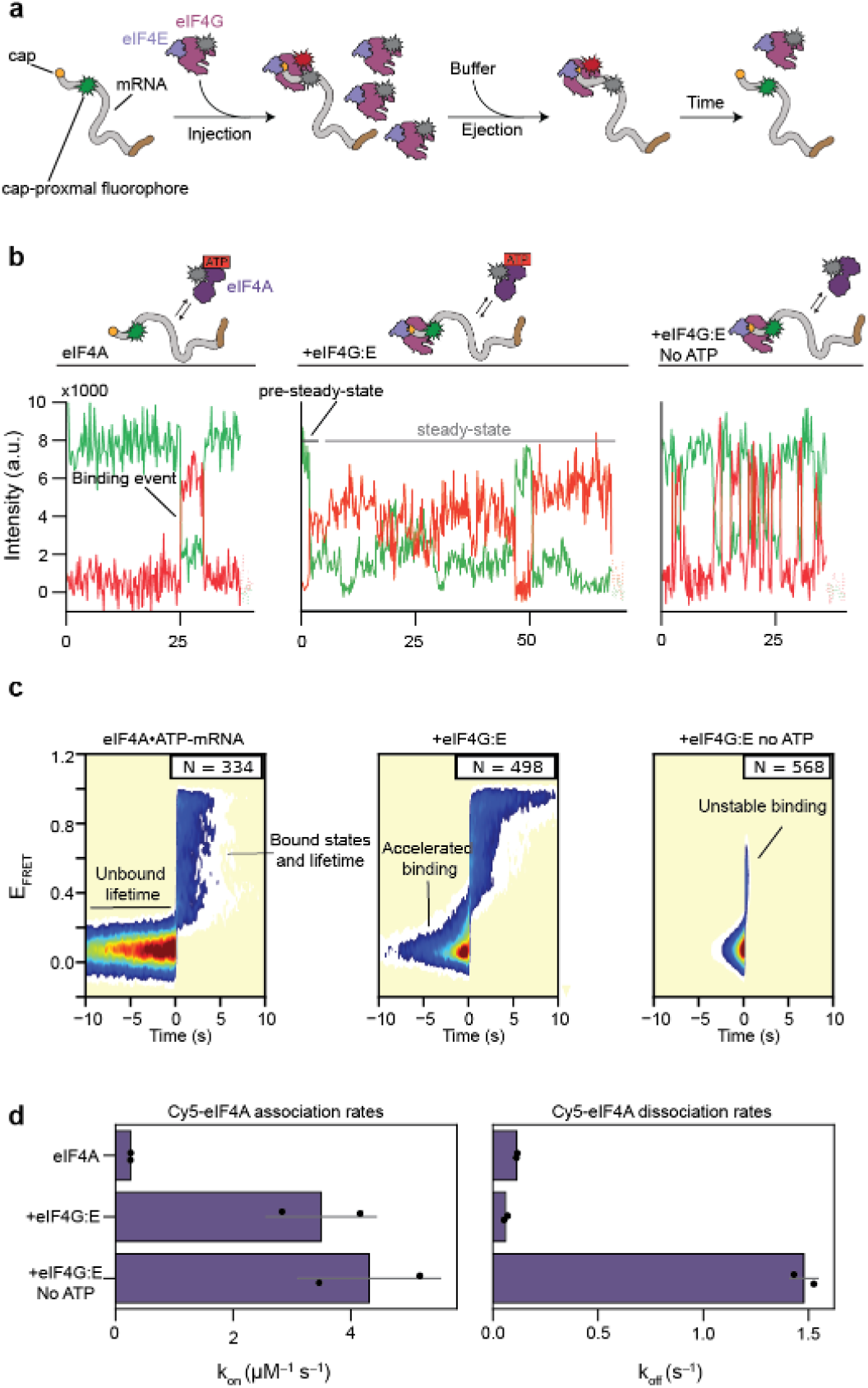
ATP binding gates productive assembly of eIF4F on mRNA. **(a)** Schematic of the single-molecule fluorescence microscopy assay used to measure pre-steady-state association and dissociation kinetics of eIF4F components on surface-tethered mRNAs in microfluidic flowcells. Fresh solutions containing initiation factors can be rapidly injected into the flowcell, whereas unbound proteins can be removed by buffer exchange to isolate dissociation kinetics. **(b)** Representative single-molecule fluorescence intensity *vs*. time trajectories showing association of Cy5-eIF4A with the capped Cy3-rpl41a mRNA construct in the absence of eIF4G:E (left), in the presence of eIF4G:E and ATP (center), or in the presence of eIF4G:E without ATP (right). Green and red trajectories correspond to Cy3 and Cy5 fluorescence intensities, respectively. **(c)** Post-synchronized surface contour plots of Cy5-eIF4A binding events measured under the conditions shown in (b). In the absence of eIF4G:E, ATP-bound Cy5-eIF4A associates slowly but remains stably bound once associated (left). Addition of eIF4G:E accelerates Cy5-eIF4A association while maintaining stable binding in the presence of ATP (center), where omission of ATP results in unstable binding despite rapid association mediated by eIF4G:E (right). **(d)** Quantification of the Cy5-eIF4A rate of association (*k*_on_) (left) and dissociation (*k*_off_) (right) measured from the experiments shown in (c). Error bars represent the standard deviation of replicate measurements, which are shown as points.

Because these initial experiments relied on ATP analogs and were performed in a time regime insensitive to kinetic defects in stripping, we sought to directly test whether ATP hydrolysis is required for eIF4A-mediated stripping of eIF4F. Furthermore, human eIF4A has been reported to hydrolyze several ATP analogs^13,15,29,30^, raising the possibility that residual ATP hydrolysis contributed to the observed stripping activity. To address this possibility, we mutated the canonical ‘DEAD’ motif in eIF4A to either ‘DEAH’ (hereafter eIF4A_slow_) or ‘DQAD’ (eIF4A_dead_), generating eIF4A constructs that either slowly hydrolyzed ATP or could not hydrolyze ATP, respectively, as measured by steady-state ATP hydrolysis assays (Extended Data Figure 1a,b)^12,31–33^. Standard mRNA recruitment gel-shift assays confirmed that eIF4F containing eIF4A_slow_ slowly stimulated cap-dependent mRNA recruitment, whereas eIF4F containing eIF4A_dead_ failed to support recruitment (Extended Data Figure 2). Having validated the expected activities of these mutants, we next tested their ability to strip Cy5-eIF4G complexed with eIF4E (Cy5-eIF4G:E) from uncapped mRNAs using smFRET (Extended Data Figure 1c). In these experiments, Cy5-eIF4G:E was first allowed to associate with the mRNA before unbound complexes were ejected and saturating eIF4A together with ATP•Mg^2+^ were injected into the flowcell. Both eIF4A_slow_ and eIF4A_dead_ stripped Cy5-eIF4G:E from the uncapped mRNAs, demonstrating that ATP hydrolysis is not required for eIF4F stripping (Extended Data Figure 1d,e).

Because ATP binding, but not hydrolysis, was required for eIF4F stripping activity, we hypothesized that ATP regulates either the composition or conformation of eIF4F on the mRNA. To distinguish between these possibilities, we site-specifically labeled eIF4A with a Cy5 acceptor fluorophore (Cy5-eIF4A) and directly monitored its binding kinetics and positioning on the mRNA. We performed pre-steady-state experiments by injecting 200 nM Cy5-eIF4A into a flowcell containing the capped mRNA construct in either the presence or absence of ATP and/or pre-bound unlabeled eIF4G:E (Figure 1b). Unexpectedly, Cy5-eIF4A•ATP associated slowly with mRNA (<0.3 µM^−1^ s^−1^, rate limited by photobleaching) but remained stably bound once associated, dissociating at only ~0.1 s^−1^, indicating that the micromolar affinity of eIF4A•ATP for mRNA arises primarily from slow association rather than rapid turnover of the bound complex (Figure 1c,d). The addition of eIF4G:E accelerated the association rate of both apo-eIF4A and eIF4A•ATP more than tenfold (~2 µM^−1^ s^−1^), whereas stable binding required ATP, with apo-eIF4A dissociating rapidly (~1.5 s^−1^) from eIF4G:E•mRNA complexes (Figure 1c,d). Together, these results demonstrate that ATP binding gates productive assembly of the ATP-bound eIF4F•mRNA complex.

Notably, association of eIF4A•ATP with mRNA-bound eIF4G:E was biphasic, likely reflecting a mixture of direct mRNA binding by Cy5-eIF4A and binding through eIF4G:E, such that the population-weighted average association rate (Fig 1c) may slightly underestimate the full effect of eIF4G:E on eIF4A recruitment (Supplemental Table 1). Moreover, transient sampling of apo-eIF4A on eIF4G:E•mRNA occupied an intermediate E_FRET_ state, suggesting that apo-eIF4A binds eIF4G:E•mRNA in a distinct conformation relative to eIF4A•ATP (Figure 1b). We also observed that eIF4A appeared to facilitate its own dissociation from eIF4G:E•mRNA complexes, as dissociation kinetics were biphasic under steady-state conditions but monophasic during pre-steady-state measurements (Extended Data Figure 3). Given the high cellular concentrations of eIF4A, this behavior would nevertheless be expected to maintain near-continuous occupancy of eIF4G:E by eIF4A *in vivo*^5^. Finally, fluorescence anisotropy measurements using fluorescein-labeled eIF4A independently validated ATP-dependent formation of the ATP-bound eIF4F•mRNA complex, revealing an eIF4G:E-dependent increase in anisotropy only in the absence of mRNA or in the presence of both mRNA and ATP (Extended Data Figure 4a,b).

Together, these results imply that productive eIF4F assembly is intrinsically coupled to the energetic state of the cell, as ATP binding regulates association of eIF4A with eIF4G:E•mRNA. These observations also reconcile an apparent distinction between lower- and higher-eukaryotic eIF4F behaviors. In both systems, ATP-binding is required to form a complete eIF4G:E:A•mRNA complex, although in the absence of ATP yeast prefers to eject eIF4A from the complex, whereas mammals appear to eject mRNA^3,13^. Moreover, these experiments indicate that ATP binding enables eIF4A to adopt a conformation capable of functioning as a strippase by displacing eIF4F from mRNA regions not stabilized by cap recognition. However, although these studies establish that ATP binding promotes productive eIF4F assembly and enables eIF4A-mediated stripping of eIF4F from mRNA, they do not identify the mechanistic role of the essential ATP hydrolysis event during translation initiation.

### The cap does not modulate ATP hydrolysis by eIF4F

Our results thus far demonstrated that ATP hydrolysis by eIF4F is not required for eIF4F stripping prior to cap recognition. We therefore reasoned that ATP hydrolysis must instead be regulated by cap recognition itself or by a downstream interaction with the PIC during mRNA recruitment. Because ATP hydrolysis by eIF4F is essential for mRNA recruitment, we considered it unlikely that ATP hydrolysis would be uncoupled from productive steps in initiation. Notably, the rate at which eIF4F is removed from the 5’ end of capped mRNAs (~0.6 min^−1^) is somewhat slower, but comparable to, the intrinsic rate of ATP hydrolysis by eIF4A on rpl41a mRNA in the absence of other initiation components (~2 min^−1^)^23^, raising the possibility that cap recognition by eIF4E allosterically suppresses both the stripping and ATP hydrolysis activities of eIF4A. In this model, productive downstream engagement with the PIC would subsequently relieve this suppression and couple ATP hydrolysis to a discrete step of translation initiation.

To test whether cap recognition regulates ATP hydrolysis by eIF4F, we performed ATP hydrolysis assays using uncapped and capped variants of a short rpl41a mRNA fragment capable of accommodating only a single eIF4F complex. The measured ATP hydrolysis rate (~10 RNA^−1^ min^−1^) was consistent with previous measurements^23,34,35^. Surprisingly, however, the eIF4F ATP hydrolysis rate was unaltered by the presence of the cap, demonstrating that cap recognition itself does not regulate ATP hydrolysis (Extended Data Figure 5). These results therefore suggest that ATP hydrolysis must instead be modulated through productive engagement with the PIC and utilized during either mRNA loading or the downstream scanning process.

### ATP hydrolysis recycles eIF4F during mRNA loading

Although eIF4F is thought to make multiple contacts with the PIC, little experimental evidence explains how eIF4F stimulates mRNA recruitment or defines the timing and mechanistic role of its essential ATP hydrolysis event^1–4,20^. Multiple models have proposed that ATP hydrolysis by eIF4F functions to displace, or recycle, the eIF4F proteins from the 5’ cap, to unwind local mRNA structure so that the PIC can load downstream of the cap, or to utilize ATP hydrolysis to facilitate scanning^1–4,9,17,36,37^. To distinguish between these possibilities, we directly monitored engagement of eIF4F with the mRNA during mRNA recruitment under conditions that recapitulate ensemble recruitment reactions (Extended Data Figure 2a). Specifically, we monitored how E_FRET_ between Cy5-eIF4G:E and the capped Cy3-rpl41a mRNA evolved during the course of a recruitment reaction. In this assay, PIC-dependent conformational changes of eIF4F are expected to alter E_FRET_ states, whereas recycling of eIF4F would abolish E_FRET_ entirely. Furthermore, by using eIF4A_slow_ and eIF4A_dead_, we reasoned that we could isolate the step at which ATP hydrolysis by eIF4F occurs.

To perform these experiments, we tethered capped Cy3-rpl41a mRNAs to the surface of the flowcell and allowed Cy5-eIF4G:E to associate with the mRNA before simultaneously ejecting unbound complexes and injecting mixtures containing varying combinations of eIF4A, ATP, and PICs composed of ribosomal 40S subunits, eIF3, eIF1, eIF1a, and the ternary complex composed of eIF2, GTP, and initiator tRNA (TC) (Figure 2a). Under these conditions, the complete PIC mixture rapidly stimulated mRNA recruitment in ensemble assays (Extended Data Figure 2a). As we previously observed^24^, Cy5-eIF4G:E remained stably associated with capped Cy3-rpl41a mRNAs for tens of minutes in the absence of additional factors, exhibiting an apparent lifetime exceeding 180 seconds (k_displace_ < 0.005 s^−1^) under imaging conditions that are effectively limited by photobleaching (Figure 2b,c)^24^. Addition of eIF4A and ATP dramatically shortened the cap residence lifetime of eIF4F to ~110 seconds (k_displace_ ~0.009 s^−1^) (Figure 2b,c). Relative to this baseline, addition of the complete PIC mixture under conditions that drive mRNA recruitment all the way to start codon recognition in ensemble assays drastically reduced the residence lifetime of eIF4F to ~25 seconds (k_displace_ ~0.04 s^−1^) (Figure 2b,c). This PIC-stimulated recycling was abolished when ATP was omitted or when eIF4A•ATP was replaced with eIF4A_slow_•ATP or eIF4A_dead_•ATP, demonstrating that productive PIC engagement stimulates ATP-hydrolysis-dependent recycling of eIF4F during mRNA loading (Figure 2c).

**Figure 2.**
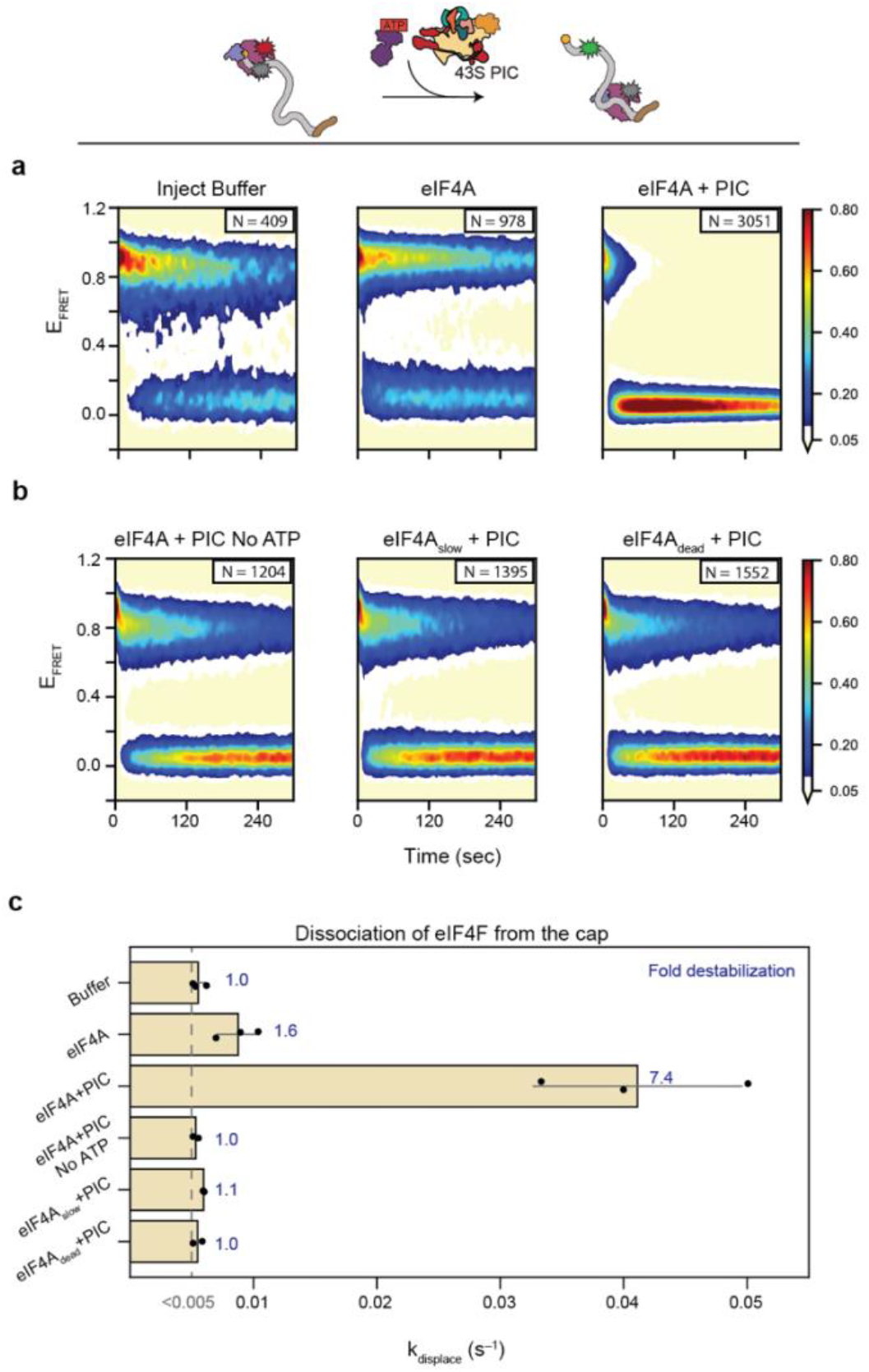
Productive PIC engagement stimulates ATP-hydrolysis-dependent recycling of cap-trapped eIF4F. **(a)** Schematic of the single-molecule fluorescence microscopy assay used to monitor recycling of cap-trapped eIF4F during mRNA recruitment. Surface-tethered capped Cy3-rpl41a mRNAs were first bound by Cy5-eIF4G:E-containing eIF4F complexes before unbound proteins were removed by buffer exchange and solutions containing varying combinations of eIF4A, ATP, and PICs were injected into the flowcell. **(b)** Surface contour plots generated using the E_FRET_ *vs*. time trajectories of cap-trapped Cy5-eIF4G:E following injection of the indicated reaction mixtures. Injection of buffer alone resulted in stable cap association of eIF4F (upper left). Addition of eIF4A and ATP modestly destabilized cap-bound eIF4F (upper center), whereas addition of both eIF4A and PICs strongly stimulated recycling of cap-trapped eIF4F (upper right). PIC-stimulated recycling was abolished in the absence of ATP (lower left) or when ATP hydrolysis-defective eIF4A mutants (eIF4A_slow_ or eIF4A_dead_) replaced wild-type eIF4A (lower center and right). **(c)** Quantification of eIF4F rate of dissociation (*k*_displace_) measured from the experiments shown in (b). Numbers indicate the fold-destabilization relative to buffer-only conditions. Error bars represent the standard deviation of replicate measurements, which are shown as points.

These results identify the essential ATP hydrolysis event during translation initiation as the step that couples PIC loading to eIF4F recycling. Moreover, our observation that eIF4F recycles from the cap implies that it must repeatedly dissociate from the cap, hop along the mRNA, and ultimately rebind the cap to catalyze each PIC-loading step in a translational burst^28^. More broadly, our results support a model in which the intrinsic rate of ATP hydrolysis by eIF4A functions analogously to a molecular timer that limits the cap residence lifetime of eIF4F, thereby releasing cap-stalled eIF4F complexes that fail to productively engage a PIC within ~2 minutes.

### eIF3 and eIF4B jointly stimulate ATP-hydrolysis-dependent recycling of eIF4F

Having established that eIF4F recycles during mRNA recruitment, we next sought to determine which PIC components stimulate this ATP-hydrolysis-dependent recycling. To do so, we repeated the recycling experiments described above while systematically omitting components from the PIC mixture (Figure 3a). Because non-core subunits of human eIF3 interact directly with eIF4F^25^ and eIF3 functionally cooperates with eIF4F in yeast^23^, we first examined the role of eIF3. In the absence of eIF3, eIF4F failed to recycle and instead behaved similarly to reactions lacking PICs altogether (Figure 3b,c), indicating that eIF3 is required to stimulate ATP-hydrolysis-dependent recycling of eIF4F.

**Figure 3.**
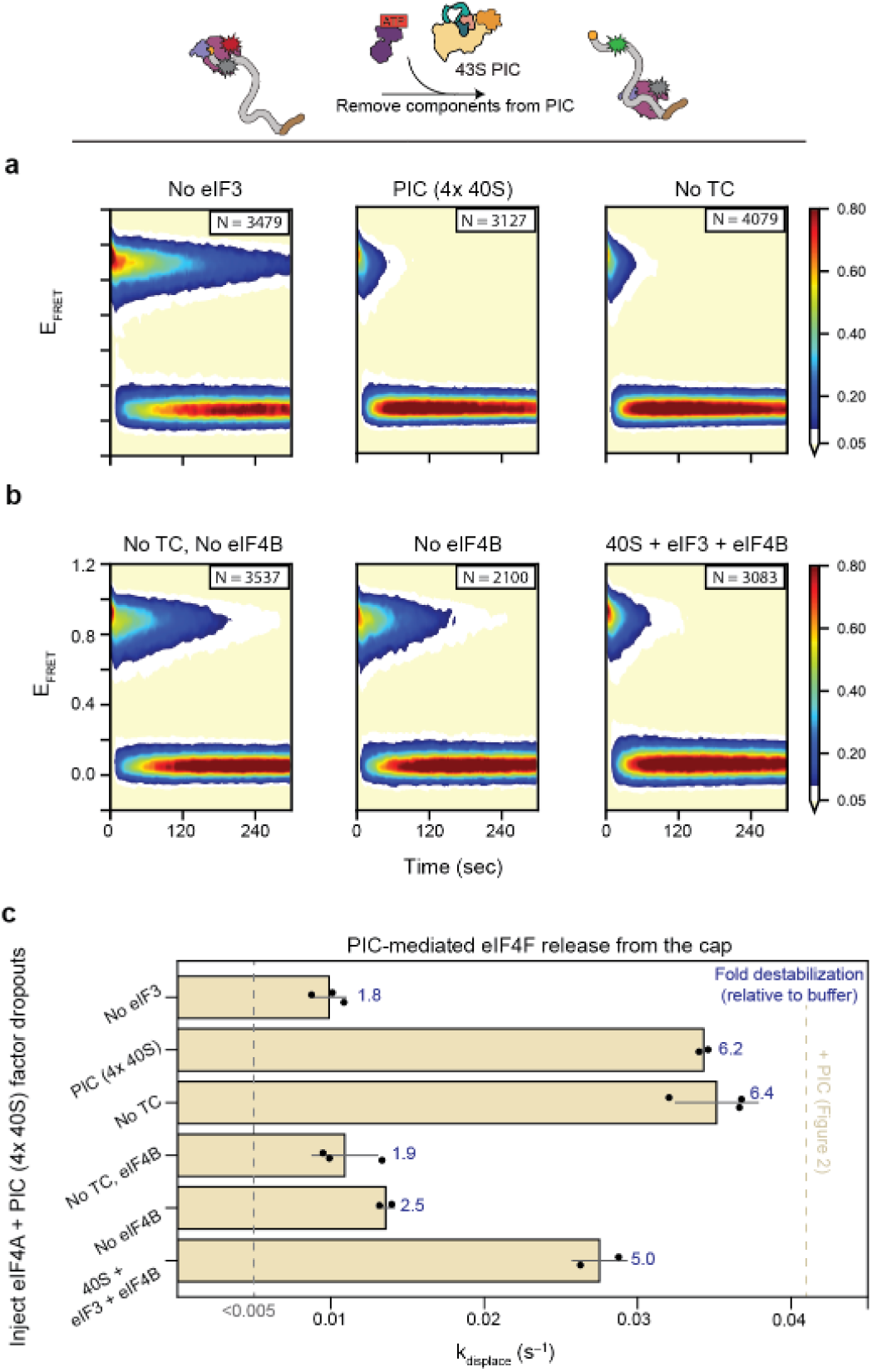
eIF3 and eIF4B jointly stimulate ATP-hydrolysis-dependent recycling of eIF4F. **(a)** Schematic of the single-molecule fluorescence microscopy assay used to monitor recycling of cap-trapped eIF4F during mRNA recruitment. Surface-tethered capped Cy3-rpl41a mRNAs were first bound by Cy5-eIF4G:E-containing eIF4F complexes before unbound proteins were removed by buffer exchange and solutions lacking individual PIC components were injected into the flowcell. **(b)** Surface contour plots of the drop-out experiments reveal that removal of eIF3 abolished efficient recycling of cap-trapped eIF4F (upper left). PIC mixtures assembled with excess 40S subunits strongly stimulated eIF4F recycling (upper center), whereas removal of TC had little effect (upper right). In contrast, removal of eIF4B strongly impaired recycling both in the presence and absence of TC (lower left and center). Efficient recycling could be partially reconstituted using only 40S subunits, eIF3, and eIF4B (lower right). **(c)** Quantification of eIF4F rate of dissociation (*k*_displace_) measured from the experiments shown in (b). Numbers indicate the fold destabilization relative to buffer-only conditions from Figure 2. Error bars represent the standard deviation of replicate measurements, which are shown as points.

To determine whether eIF3 functions in this capacity as part of the 43S PIC, we repeated these experiments in the absence of 40S subunits (Extended Data Figure 6a,b). Surprisingly, mixtures lacking 40S subunits stimulated eIF4F recycling with little defect (Extended Data Figure 6b,c), raising the possibility that free eIF3 present in excess over 40S subunits in the PIC mixture mediates this activity. Consistent with this interpretation, free eIF3 stimulated eIF4F recycling much less effectively than complete PICs, whereas sequestration of eIF3 by excess 40S subunits abolished this activity (Extended Data Figure 6b,c). Importantly, however, full recycling activity was restored when complete 43S PICs were assembled in the presence of excess 40S subunits while maintaining constant concentrations of the remaining initiation factors (Figure 3b). Together, these results indicate that PIC-bound eIF3 requires at least one additional initiation factor to stimulate eIF4F recycling during productive PIC engagement. Moreover, the inability of 40S subunit-bound eIF3 alone to efficiently stimulate recycling suggests that the 40S subunit, which is present in large excess *in vivo*^5,38,39^, may constrain eIF3 in a conformation that suppresses non-productive eIF4F recycling.

To identify the additional factor required for productive eIF4F recycling, we systematically removed individual initiation factors from PIC mixtures assembled with elevated concentrations of 40S subunits. Unexpectedly, removal of TC had little effect on eIF4F recycling, whereas removal of eIF4B abolished the ability of 40S subunit-bound eIF3 to stimulate recycling (Figure 3b,c). Moreover, efficient recycling could be reconstituted using only 40S subunits, eIF3, and eIF4B (Figure 3b,c), indicating that eIF4F recycling does not require a fully assembled PIC. Although the precise role of eIF4B in translation initiation has remained unclear, its functional cooperation with both eIF3 and eIF4F is consistent with known interactions between eIF4B and both complexes^22^, as well as previous observations that eIF4B does not directly contribute to cap recognition^24^. Our results further suggest that eIF4B functions together with PIC-bound eIF3 to stimulate the ATP hydrolysis event required for eIF4F recycling, consistent with observations that eIF4B can associate directly with the 40S subunit^27,40^.

Finally, we sought to validate the synergy between eIF3, eIF4B, and eIF4F using ensemble mRNA recruitment assays. As expected, standard mRNA recruitment reactions using capped rpl41a mRNAs showed that recruitment remained slow in the absence of eIF4B (Extended Data Figure 2b,c). To more directly test functional synergy between eIF4F and eIF3, we next examined recruitment of a capped poly(CAA) model mRNA (Extended Data Figure 7a-d). Unlike native mRNAs such as *rpl41a*, recruitment of this model mRNA is stimulated, but not strictly dependent upon, either eIF4F or eIF3, thereby allowing factor drop-out experiments to assess functional synergy^41,42^. We therefore performed recruitment reactions containing fixed concentrations of eIF4G:E, eIF1, eIF4B, eIF1A, 40S subunits, and TC while titrating eIF4A in either the presence or absence of eIF3. Under these conditions, efficient stalling of eIF4G:E at the cap requires eIF4A, thereby minimizing non-specific association of eIF4G:E near the cap-proximal position on the mRNA. In the absence of eIF3, cap-stalled eIF4F inhibited recruitment of the PIC to the model mRNA, indicating that eIF4F was unable to recycle from the cap. Upon addition of eIF3, however, this inhibition was converted into stimulation, and this stimulation remained dependent upon eIF4B, consistent with our single-molecule experiments (Extended Data Figure 7).

Together, these results demonstrate that eIF3 and eIF4B jointly stimulate ATP-hydrolysis-dependent recycling of eIF4F during mRNA loading. Notably, the maximal rate of eIF4F recycling we measured here closely matches the maximal rate of mRNA recruitment observed here and previously *in vitro*^22–24^, strongly suggesting that ATP-hydrolysis-dependent recycling of eIF4F and productive PIC loading are mechanistically coupled during mRNA recruitment. Under conditions where initiation factors are abundant, this coupled recycling/loading step therefore likely constitutes the rate-limiting step of mRNA recruitment. These observations further suggest that productive PIC loading and ATP-hydrolysis-dependent eIF4F recycling occur concomitantly under ideal conditions. Furthermore, the observation that eIF4F recycling does not require eIF1, eIF1A, or TC suggests either that eIF4F can become uncoupled from fully productive PIC loading under some conditions or that partially assembled PICs can engage mRNAs prior to complete maturation. In the latter model, PIC maturation could therefore continue after PIC loading and potentially during downstream scanning.

## Discussion

The mechanism by which a cap-stalled eIF4F facilitates PIC loading has remained unresolved since the discovery of eIF4F. Early models proposed that eIF4F might use ATP hydrolysis to recycle itself or other cap-associated proteins during PIC loading, thereby allowing the PIC to occupy the vacated site^12^. More recent models instead propose that cap-stalled eIF4F remains positioned at the 5’ end of the mRNA and uses ATP hydrolysis to remodel local RNA structure, thereby enabling the PIC to load downstream of eIF4F^1–4,9,17,36,37^. Additional models have further proposed that cap-stalled eIF4F remains bound to the scanning PIC or that ATP hydrolysis by eIF4F, or by eIF4A together with eIF4B, directly drives PIC scanning through translocase-like activity^9,21^.

Contrary to current models, our single-molecule experiments support a mechanism in which the essential ATP hydrolysis step catalyzed by eIF4F promotes recycling of cap-stalled eIF4F during productive PIC engagement. We show that ATP binding, but not ATP hydrolysis, is required for stable association of eIF4A with mRNA-bound eIF4G:E and that the resulting ATP-bound eIF4F complex is continuously displaced from uncapped positions on the mRNA until it stochastically samples and becomes kinetically trapped at the cap. ATP hydrolysis instead promotes destabilization and recycling of the cap-stalled eIF4F complex. Our results indicate that ATP-hydrolysis-dependent recycling of eIF4F occurs prior to scanning and during the PIC loading phase of translation initiation, consistent with previous ensemble and single-molecule studies of mRNA recruitment and scanning^20,23^. Moreover, we show that eIF3 and eIF4B cooperate to stimulate ATP-hydrolysis-dependent recycling of eIF4F, thereby functionally coupling eIF4F recycling to productive PIC engagement. Finally, we speculate that the large binding footprint of eIF4F on the mRNA^24^ may transiently maintain the mRNA in a loading-competent conformation that facilitates exchange of eIF4F for the PIC during loading, analogous to how unstructured mRNA sequences can promote eIF4F-independent loading^22,41^.

Together with our previous work^24^, these results position eIF4A as a central regulator of eIF4F positioning and dynamics on mRNA and further suggest that the intrinsically slow ATP hydrolysis rate of eIF4A functions analogously to a timer that limits the amount of time available for productive PIC engagement with cap-stalled eIF4F, preventing the limited pool of eIF4F from being trapped on mRNAs unable to productively engage with PICs. Our results further imply that eIF4A occupies multiple conformational states within eIF4F that have yet to be structurally visualized^34,43–46^. In addition, these observations define a role for eIF4B, whose function in translation initiation has remained incompletely understood, in assisting eIF3-dependent stimulation of ATP-hydrolysis-dependent eIF4F recycling, potentially through regulation of eIF4G conformational dynamics. Finally, the functional cooperation we observe between eIF3 and eIF4G further suggests that these factors interact directly in yeast despite the absence of the C-terminal eIF3-binding extension found in mammalian eIF4G, perhaps through a short motif within the core eIF3c subunit of eIF3^3,25^.

Unexpectedly, we observed efficient eIF3-driven eIF4F recycling in the absence of eIF1, eIF1A, and TC, suggesting that complete PIC assembly is not strictly required for eIF4F recycling and PIC loading. These observations raise two non-mutually exclusive possibilities: partially assembled PICs may undergo non-productive loading events, or incompletely assembled PICs may engage and load onto mRNAs prior to full PIC assembly, with remaining assembly steps occurring before or during downstream scanning and/or start codon selection^47^. Although PICs likely exist in multiple compositional states with varying loading efficiencies, we favor the latter possibility, at least in the case of TC, because loading of TC-deficient PICs could permit scanning under conditions of limiting TC availability and thereby facilitate bypassing of upstream open reading frames (uORFs) without requiring re-initiation. This interpretation is consistent with current models of re-initiation, which already invoke TC-deficient scanning complexes, as well as *in vitro* biochemical observations that mRNA accelerates TC association with PICs^*48*^.

Collectively, our results (Extended Data Table 1) support a new model for eIF4F-dependent mRNA recruitment (Figure 4). In the first step, eIF4F stochastically samples the mRNA, with eIF4A-driven stripping facilitating its sampling until it becomes transiently trapped at the cap. During this interval, the intrinsically slow ATP hydrolysis rate of eIF4A functions analogously to a timer, limiting the availability of eIF4F for productive PIC engagement before it recycles and potentially samples a competing mRNA. Upon encountering a PIC, eIF3 and eIF4B jointly stimulate ATP hydrolysis by eIF4F, reactivating the strippase modality of eIF4A and triggering recycling of eIF4F from the cap, which forces eIF4F to hop to a different location on the mRNA or to dissociate entirely. We speculate that 40S subunit loading occurs concomitantly with eIF4F release and that the large footprint of eIF4F on the mRNA transiently maintains the mRNA in a loading-competent conformation, thereby enabling the 40S subunit to occupy the space previously bound by eIF4F. Simultaneously, ATP hydrolysis by eIF4F may also induce eIF3-mediated conformational rearrangements of the PIC that help drive PIC loading, consistent with proposals from mammalian systems^15^. Following this exchange, any missing initiation factors (*e*.*g*., TC) could subsequently associate with the PIC before loading has completed^49^ and/or during downstream scanning.

**Figure 4.**
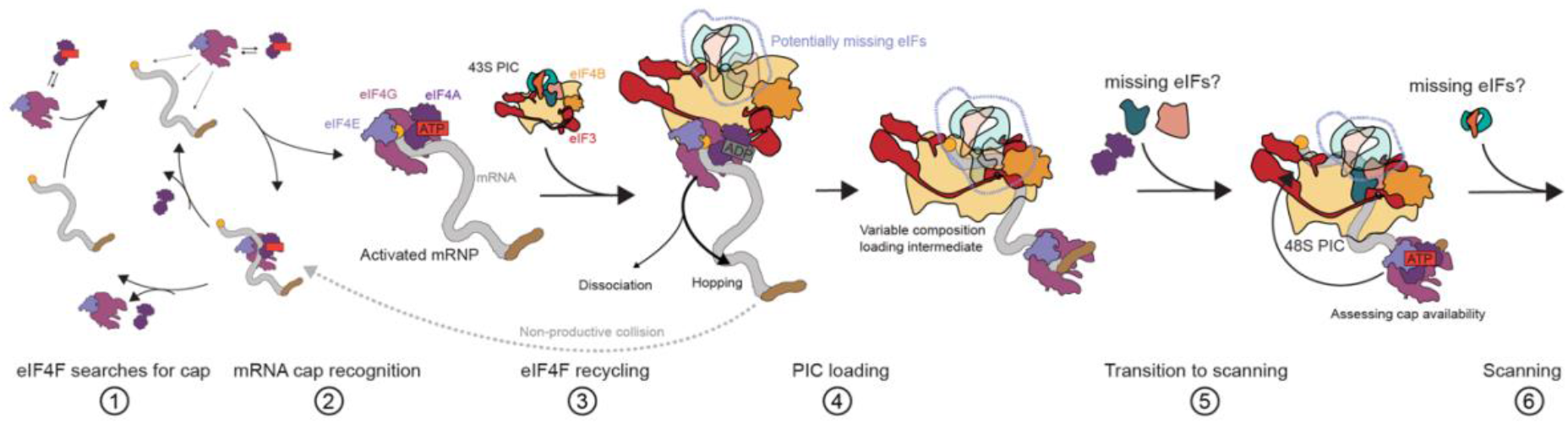
Mechanistic model for eIF4F-dependent cap recognition and PIC loading during mRNA recruitment. Step 1. eIF4F dynamically redistributes along the mRNA through ATP-dependent, eIF4A-mediated stripping from cap-distal positions. Step 2. Upon recognition of the 5′ cap by eIF4E, eIF4F becomes transiently trapped at the cap, generating an activated messenger ribonucleoprotein (mRNP) competent for productive PIC engagement. During this interval, the intrinsically slow ATP hydrolysis rate of eIF4A functions as a molecular timer, limiting the time available for productive PIC engagement before eIF4F recycles. Step 3. Productive engagement of cap-trapped eIF4F by the PIC stimulates ATP-hydrolysis-dependent recycling of eIF4F through the combined activities of eIF3 and eIF4B. Recycled eIF4F can either hop to another position along the mRNA or dissociate from the transcript to engage a different mRNA, thereby enabling continued redistribution. Non-productive PIC encounters fail to trigger productive PIC loading before eIF4F recycling occurs. Step 4. ATP-hydrolysis-dependent recycling of eIF4F is coupled to loading of the PIC onto the mRNA. We speculate that the large footprint of eIF4F may transiently maintain the mRNA in a loading-competent conformation that facilitates this exchange. Step 5. Following PIC loading, initiation factors absent from the incipient PIC can associate before and/or during downstream scanning, ultimately yielding a scanning 48S PIC (Step 6) competent for start codon recognition. Because our data suggest that productive PIC loading can occur with incompletely assembled PICs, the timing of incorporation of individual initiation factors may vary and could occur at any point between steps 3 and 6. Consistent with biochemical evidence suggesting that eIF1 and/or eIF1A are required for scanning^47^, the latest possible incorporation of these factors is depicted at the transition from loading to scanning, whereas TC is shown associating during scanning to illustrate a potential mechanism for facilitating uORF bypass under conditions of limiting TC availability.

Importantly, the ability of eIF4F to hop^24^ to another location on the mRNA after recycling from the cap would enable a single copy of eIF4F to remain associated with the transcript and potentially rebind the cap after the 48S PIC scans downstream. This retention could allow a single eIF4F complex to drive multiple rounds of initiation, consistent with expectations from *in vivo* single-molecule imaging^28^ while eliminating the need to invoke cap-tethered scanning^21^ and still permitting the initial recruitment of eIF4F to remain rate-limiting^24^. Furthermore, because the 5’ and 3’ ends of mRNAs are often in close physical proximity^50,51^ and poly(A) binding protein, located at the 3’ end, directly interacts with eIF4G^52^, we speculate that recycled eIF4F may preferentially redistribute toward the 3’ end of the mRNA after recycling. In this scenario, because increasing ribosome density is known to progressively separate the mRNA ends^53^, each productive initiation event would reduce the probability that eIF4F remains associated with the mRNA after successive rounds of recycling and ultimately limit the maximal ribosome density on a given mRNA^50,54^. More broadly, our model suggests that cap-dependent translation initiation can be regulated at two distinct levels: factors that promote initial acquisition of eIF4F by an mRNA and factors that promote retention of eIF4F between successive rounds of PIC loading. In both cases, the critical mechanistic feature is the ability of eIF4F to dynamically redistribute along the mRNA, suggesting that mRNA structural elements and RNA-binding proteins that modulate mRNA topology may ultimately exert broad control over cap-dependent translational efficiency.

## Materials and Methods

### Protein expression and purification

The system used to generate the mRNAs, tRNAs, ribosomes, and proteins employed in this study has been described previously^24^. In brief, yeast ribosomes were isolated from the *S. cerevisiae* strain W303 following established methods^55^. The yeast initiation factors eIF1, eIF1A, eIF4A, eIF4G:E, and eIF4B were expressed and purified from *Escherichia coli* (*E*. Coli) BL21 RIPL cells (Agilent) as previously described^24^. eIF3 was expressed and purified from *E. coli* BL21 RIPL cells as the individual subunits eIF3a, eIF3b, eIF3c, and eIF3i-g and subsequently reconstituted into a functional pentameric complex as previously described^42,56,57^.

### Protein labeling

To enable site-specific fluorophore labeling while minimizing background labeling of endogenous cysteines, the mutations C250A and T189C were introduced into eIF4A by site-directed mutagenesis. eIF4A(C250A/T189C) was purified identically to wild-type eIF4A and stored in the presence of a 4-fold molar excess of tris(2-carboxyethyl)phosphine (TCEP) as the sole reducing agent.

For fluorophore labeling, eIF4A(C250A/T189C) was incubated overnight at 4 ◦C with either a 3-fold molar excess of fluorescein (FAM)-maleimide (Lumiprobe) or sulfo-Cy5-maleimide (Lumiprobe) in the presence of 1% DMSO. Unreacted fluorophores were removed by passing the reaction mixture through 5 5-mL desalting columns (Cytiva) connected in series. The labeled protein was subsequently purified by gel filtration using a Superdex 75 10/300 GL column (Cytiva). Proteins were exchanged into storage buffer containing 20 mM 2-[4-(2-hydroxyethyl)piperazin-1-yl]ethanesulfonic acid (HEPES pH = 7.4), 100 mM potassium acetate (KOAc), and 10% glycerol, flash frozen in single-use aliquots, and stored at –80 ºC.

Cy5-eIF4G:E was generated by reacting eIF4G(501C):E overnight with a 10-fold molar excess of sulfo-Cy5-maleimide, followed by purification by gel filtration and mono S chromatography as previously described^24^.

### RNA preparation

The initiator tRNA used for mRNA recruitment assays was *in vitro* transcribed, purified, and aminoacylated as previously described^24,55^. All single-molecule experiments were performed using an mRNA derived from the *S. cerevisiae rpl41a* gene containing a 24-nucleotide 5’ untranslated region (UTR). The rpl41a mRNA was site-specifically fluorophore-labeled at position A14 using a terbium-assisted deoxyribozyme^58^ as previously described^24^. Where indicated, rpl41a mRNA was capped to homogeneity using the Vaccinia Capping System (New England Biolabs).

For single-molecule experiments, rpl41a mRNA was tethered to the flowcell surface through hybridization to a biotinylated DNA oligonucleotide complementary to an internal region of the transcript. The sequences of all the RNAs and DNA oligonucleotides used in this study are listed below.

rpl41a mRNA sequence: GGAGACCACAUCGAUUCAAUCGAAAUGAGAGCCAAGUGGAGAAAGAAGAGAACUAGAA GACUUAAGAGAAAGAGACGGAAGGUGAGAGCCAGAUCCAAAUAAGCGGAUUAUGAGUA AAUAACUCUAAUUUUGUUUUAAAUUCUUUCAAGAGUAUCGUAAUGUCAUUGAUGAAUUA ACAUGUUAGUUUCUAUUCUACCUCAUAAUGGAUCUAAAUUGCAUACUAAUCUCACGGU GGGGUGUAAACCAUUGCCUACUAUUUAUAUAGUGCUUUAUAUAUGUCUCACAUAGUUU AAUCAAUUGUCCGUUUUUUUGAAAAAAAAAAAAAAAAAAAAAAAAAAAAAA

First 45 nucleotides of rpl41a mRNA: GGAGACCACAUCGAUUCAAUCGAAAUGAGAGCCAAGUGGAGAAAG

Poly(CAA) model mRNA: GGCAACAACAACAACAACAACAAAUGGAACAACAACAACAAC

Surface tethering DNA oligonucleotide (Integrated DNA technologies): Biotin-ACATGTTAATTCATCAATGAC

A14-labeling DNA oligonucleotide (Integrated DNA technologies): CCGTCGCCATCTCCCGTAGGTGAAGGGCTTGAAGGTTCCATTCCCGATGTGGTCTC

### Reaction buffer

Unless otherwise indicated, all experiments were performed in Reaction Buffer containing 20 mM HEPES (pH = 7.4), 100 mM KOAc, 3 mM magnesium acetate (Mg(OAc)_2_)^24,55,59^.

### Fluorescence anisotropy

Fluorescence anisotropy titrations were performed in Reaction Buffer using 20 nM FAM-eIF4A and a 10-point, 2-fold dilution series of eIF4G or eIF4G:E spanning concentrations from 0 to 1,727 nM. Reactions were incubated for 30 minutes at room temperature prior to measurement.

Endpoint anisotropy assays were performed in Reaction Buffer containing 20 nM FAM-eIF4A together with the following components, as indicated in the figure: 600 nM rpl41a mRNA lacking a poly(A) tail, 400 nM eIF4G:E, 2 mM ATP•Mg, and/or 500 ng nuclease A.

### NADH-coupled ATPase assays

NADH-coupled assays were performed in 96-well plates using 200 µL reaction volumes containing 100 nM RNA, 4% Lactate dehydrogenase-pyruvate kinase mixture (Sigma-Aldrich P0294), 500 µM phosphoenolpyruvate, 100 µM NADH, 2 mM ATP•Mg, 2 µM eIF4A, and 300 nM eIF4G or eIF4G:E. The RNA substrate, eIF4A variant, and eIF4G-containing complex used in each experiment are indicated in the corresponding figure panels. Experiments performed using the 45-nucleotide RNA construct contained 100 nM RNA and 50 nM eIF4G:E.

Reactions were initiated by the addition of ATP and monitored at a wavelength of 340 nm in a Biotek plate reader at 20-second intervals for 3 hours. The linear portion of each progress curve was fit by linear regression, and ATP hydrolysis rates were calculated from the rate of NADH consumption assuming a 1:1 stoichiometry between NADH oxidation and ATP hydrolysis^60^. At a reaction volume of 200 µL, the effective optical pathlength of each well was approximately 0.5 cm.

### Single-molecule fluorescence microscopy

The laboratory-built, prism-based total internal reflection fluorescence (TIRF) microscopy system and general experimental procedures used in this study have been described previously^24^. Briefly, samples were illuminated with a 20 mW laser prior to prism excitation, and images were acquired using a 200-ms camera exposure time. Experiments were performed either under continuous illumination or with the excitation laser shuttered every 3 seconds to extend fluorophore lifetimes. These acquisition modes enabled measurements on either second or minute timescales and are indicated below where appropriate.

Stripping experiments using uncapped rpl41a mRNA (Extended Data Figure 1) were performed under continuous illumination. For these experiments, Cy5-eIF4G:E was first allowed to associate with uncapped rpl41a mRNA containing a cap-proximal fluorophore. eIF4A and ATP•Mg were then injected into the flowcell as previously described^24^. However, rather than wild-type eIF4A, these experiments were performed using either eIF4A_slow_ or eIF4A_dead_.

Experiments examining association of Cy5-eIF4A with mRNA were performed by injecting 200 nM Cy5-eIF4A onto tethered, capped Cy3-rpl41a mRNA. The injection solution contained either 2 mM ATP•Mg or no ATP, as indicated in the figure panels. Where indicated, the mRNA was pre-incubated with 3 nM unlabeled eIF4G:E prior to Cy5-eIF4A injection. Ejection experiments were performed by first allowing Cy5-eIF4A to associate with the mRNA and then removing unbound protein by injecting a solution lacking eIF4A but containing ATP•Mg at the start of image acquisition.

Experiments examining ATP-hydrolysis-dependent recycling of cap-stalled eIF4G:E were performed using laser shuttering every 3 seconds. This acquisition scheme renders spontaneous dissociation of cap-stalled eIF4G:E effectively photobleaching-limited, as previously measured^24^, while allowing eIF4A-dependent recycling of eIF4G:E to occur on a timescale modestly faster than fluorophore photobleaching^24^. Consequently, these conditions provide a sensitive assay for factors that accelerate recycling of cap-stalled eIF4F.

For these experiments, 3 nM Cy5-eIF4G:E was first allowed to associate with the cap of tethered, cap-proximally labeled Cy3-rpl41a mRNA. Following association, mixtures containing PIC components together with eIF4A and ATP•Mg were injected into the flowcell. To assemble PIC mixtures, a 2× stock was prepared by first forming 3.5 µM eIF2•GTP•Met-tRNAi ternary complex (TC) and subsequently combining it with additional initiation factors to yield final concentrations of 2 µM eIF1, 2 µM eIF1A, 600 nM TC, 400 nM eIF3, 200 nM 40S subunits, and 800 nM eIF4B. This 2× mixture was diluted two-fold into microscope imaging buffer containing an oxygen-scavenging system and triplet-state quenchers^24^. ATP•Mg and eIF4A were then added to final concentrations of 2 mM and 2 µM, respectively, immediately prior to injection into the flowcell. For shuttered-acquisition experiments, injections were performed between frames 2 and 3 of each movie.

### Analysis of smFRET data

All analyses of smFRET data were performed using custom-written, publicly available software developed in our laboratory. Individual Cy3 and Cy5 fluorescence intensity *vs*. time trajectories were extracted from movies using vbSCOPE and subsequently analyzed using tMAVEN^61,62^.

For each data point in each pair of Cy3 and Cy5 fluorescence intensity *vs*. time trajectories (intensity trajectories), E_FRET_ was calculated as the Cy5 fluorescence intensity divided by the sum of the Cy3 and Cy5 fluorescence intensities and E_FRET_ *vs*. time trajectories (E_FRET_ trajectories) were calculated. Signal-to-background ratios were then calculated on an intensity trajectory-by-trajectory basis, and trajectories with signal-to-background ratios less than 3 for Cy5-eIF4A or less than 5 for Cy5-eIF4G:E were excluded from further analysis. The remaining intensity trajectories were visually inspected for characteristics consistent with single fluorophores, including single-step photobleaching or photoblinking, rather than multistep bleaching indicative of multiple molecules. After discarding intensity trajectories containing multiple molecules, the photobleaching point was manually assigned for each trajectory. Notably, in experiments performed with laser shuttering, not all trajectories were photobleached during acquisition. In such cases, intensity trajectories displaying photophysical and intensity characteristics similar to those of confirmed single-molecule trajectories were retained for analysis.

Following the identification of intensity trajectories arising from single molecules, the corresponding E_FRET_ trajectories were idealized using a three-state hidden Markov model implemented through the vbFRET algorithm. The resulting Viterbi paths were clustered using a k-means algorithm, yielding states with E_FRET_ values near 0.05, 0.40, and 0.80. For rate calculations, the two higher-E_FRET_ states (0.40 and 0.80) were combined into a single bound state. This approach is equivalent to using a 3-state model and thresholding to find the bound state with a cutoff of E_FRET_ > 0.2. Cy5-eIF4A experiments performed in the absence of ATP were instead analyzed using a two-state model.

All kinetic rates were determined using survival analysis of dwell-time distributions. Pre-steady-state association rates were calculated from dwell-time distributions compiled from the interval between Cy5-eIF4A injection and the first binding event. Pre-steady-state dissociation rates were similarly calculated from dwell-time distributions compiled from the interval between ejection of a pre-bound molecule (Cy5-eIF4A or Cy5-eIF4G:E) and the first dissociation event. Steady-state association and dissociation rates were calculated from dwell-time distributions compiled from all binding or dissociation events, respectively, excluding the first and last event in each E_FRET_ trajectory.

Experiments performed using laser shuttering, which assayed the residence lifetime of cap-stalled Cy5-eIF4G:E, focused exclusively on pre-steady-state dissociation kinetics.

Survival curves from pre-steady-state experiments were fit to single-exponential decay functions (Equation 1), whereas survival curves from steady-state experiments were fit to bi-exponential decay functions (Equation 2). For graphical presentation, population-weighted average rates derived from the bi-exponential fits (Equation 3) are reported to facilitate comparison with pre-steady-state rates. The complete fitting parameters and population weights are provided in Supplementary Table 1.

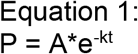

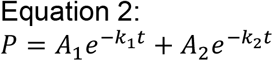

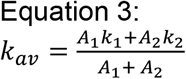

Surface contour plots were generated using tMAVEN and provide a visualization of the ensemble population distribution reconstructed from individual E_FRET_ trajectories. These plots were generated, following field standards, by lightly smoothing (smoothing parameter = 0.05 in tMAVEN) in the x- and y-directions and interpolating (parameter = 800 in tMAVEN) the data.

### mRNA recruitment gel-shift assays

Standard mRNA recruitment assays using hot-capped rpl41a mRNA were performed as previously described^24^, with the indicated substitution of eIF4A mutants or omission of eIF4B.

For recruitment assays using the poly(CAA) model mRNA, the transcript was hot-capped as previously described^24^. Subsequently, 2× PIC mixtures were assembled following the same procedure used for the microscopy experiments described above, containing final concentrations of 2 µM eIF1, 2 µM eIF1A, 600 nM TC, 100 nM eIF4G:E, 100 nM 40S subunits, and 800 nM eIF4B. These 2× mixtures were diluted into Reaction Buffer supplemented with the indicated concentration of eIF4A, 2 mM ATP•Mg, and 200 nM eIF3, when included.

Reactions were initiated by the addition of hot-capped mRNA to a final concentration of 15 nM. Reactions were incubated for 1 hour at 26 °C and then quenched by loading onto a running native 4% (37.5:1 acrylamide:bisacrylamide) tris–HEPES–EDTA–magnesium (THEM) gel. Gels were exposed to a phosphor screen and imaged using a Typhoon 5 scanner (Cytiva).

eIF4A was titrated at concentrations of 0, 100, 500, 1,000, and 5,000 nM.

### Statistical Analyses

All single-molecule experiments were performed as two or three biological replicates and nearly every individual single-molecule experiment contained >100 trajectories from which survival probabilities could be determined. All reported rates are the means determined from the set of replicates and the reported errors are the standard deviations associated with those means. The raw data points used to determine the means and standard deviations are plotted on each bar plot. The number of trajectories found in each replicate, along with replicate numbers and all fitting parameters can be found in **Supplemental Table 1**.

## Data availability

All data supporting the findings of this study are available within the manuscript and its Extended Data Figures. The number of molecules analyzed and the raw survival-curve fitting parameters for each replicate, prior to correction for frame rate and labeling efficiency, are provided in Supplementary Table 1. Intensity trajectories extracted using vbSCOPE for each experiment prior to filtering, together with the fully processed datasets used for analysis, are available through Zenodo (DOI: 10.5281/zenodo.20672937). Raw movie files are available from R.L.G. upon reasonable request.

## Code availability

The software used for analysis of single-molecule data is freely available through the Gonzalez Biophysics Lab GitHub:

### vbSCOPE

https://github.com/GonzalezBiophysicsLab/vbscope-paper

### tMAVEN

https://github.com/GonzalezBiophysicsLab/tmaven

## Acknowledgements

This work was supported by funds to R.L.G from the NIH (R01 CA277727, R01 GM 084288, and R35 GM153724). R.C.G was supported by a postdoctoral fellowship from the Helen Hay Whitney Foundation and N.A.I was supported by an NSF GRFP fellowship (DGE 1644869) and a postdoctoral fellowship from the Helen Hay Whitney Foundation. Access to the Horiba Fluorimeter in the Columbia University Precision Biomolecular Characterization Facility (PBCF) was enabled through NSF award 1828491. The authors would like to thank the Columbia University PBCF for the maintenance of and access to the equipment.

## Author contributions

This work was conceptualized by R.L.G, R.C.G, and N.A.I. The single-molecule fluorescence microscopy experiments were performed, analyzed, and interpreted by R.C.G with input from R.L.G, C.K.T, and N.A.I.. The mRNA recruitment gel-shift experiments were performed, analyzed, and interpreted by N.A.I. with input from R.L.G, R.C.G and C.E.A. R.C.G and N.A.I. generated the materials with contributions from V.M.C. R.L.G and R.C.G. wrote the paper.

---

**Extended Data Figure 1.**
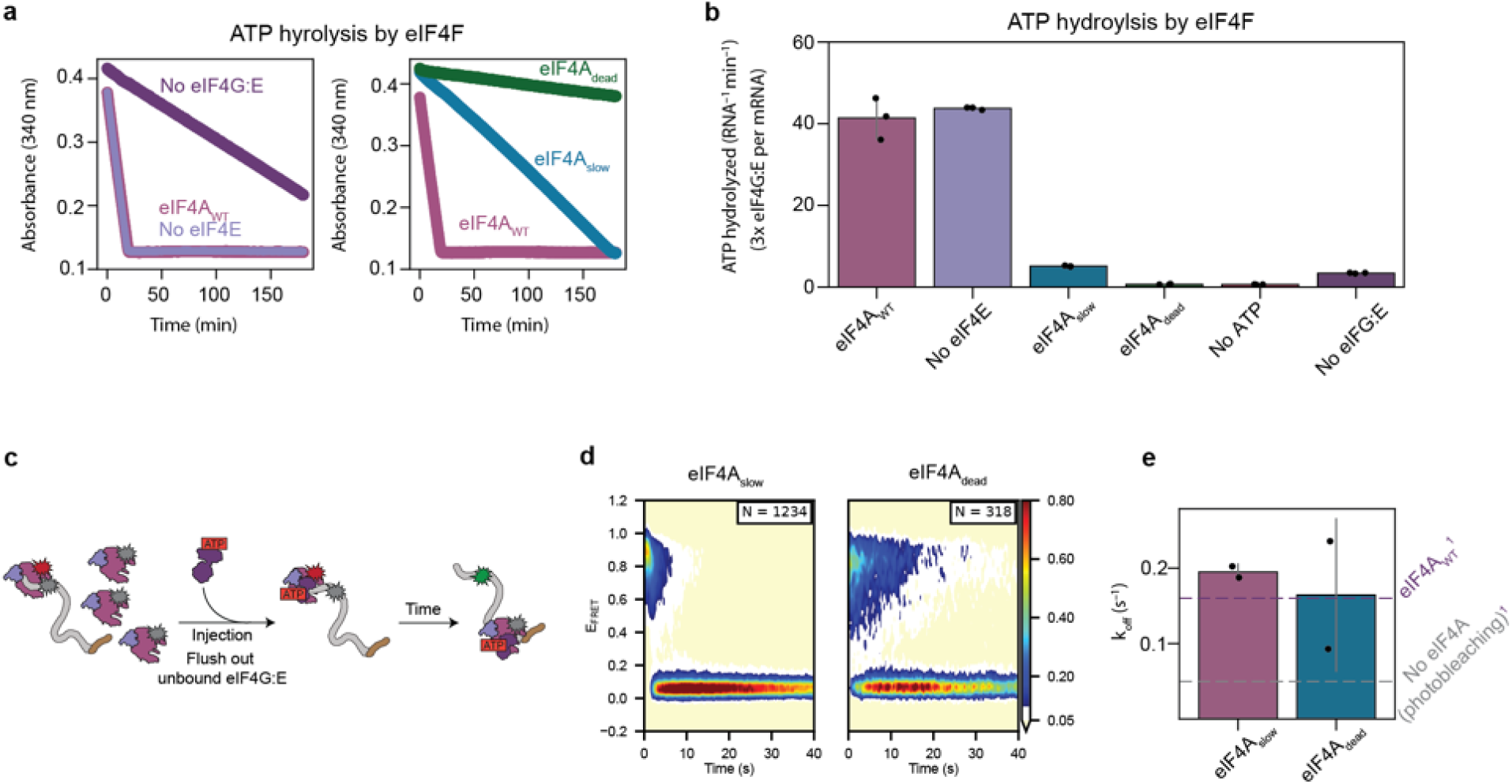
ATP hydrolysis by eIF4A is not required for displacement of eIF4G:E from uncapped mRNA. **(a)** Representative NADH-coupled ATPase assays monitoring ATP hydrolysis by eIF4A in the presence or absence of eIF4G:E (left) and by wild-type eIF4A, eIF4A_slow_, and eIF4A_dead_ in the presence of eIF4G:E (right). ATP hydrolysis was monitored through loss of NADH absorbance at 340 nm. **(b**) Quantification of ATP hydrolysis rates from assays shown in (a). eIF4G:E stimulates ATP hydrolysis by wild-type eIF4A, whereas ATP hydrolysis is strongly reduced in eIF4A_slow_ and nearly abolished in eIF4A_dead_. Error bars represent the standard deviations of replicate measurements, which are shown as points. **(c)** Schematic of the single-molecule assay used to measure displacement of Cy5-eIF4G:E from uncapped rpl41a mRNA. Cy5-eIF4G:E was first allowed to associate with uncapped mRNA, unbound complexes were removed by buffer exchange, and eIF4A mutants together with ATP•Mg were subsequently introduced into the flowcell. **(d)** Surface contour plots showing dissociation of Cy5-eIF4G:E from uncapped mRNA following addition of eIF4A_slow_ or eIF4A_dead_. Loss of the high-E_FRET_ population corresponds to displacement of eIF4G:E from the mRNA. **(e)** Quantification of eIF4G:E dissociation rates measured in (d). Both eIF4A_slow_ and eIF4A_dead_ promote displacement of eIF4G:E from uncapped mRNA despite severely compromised ATP hydrolysis activity, demonstrating that ATP hydrolysis is not required for eIF4A-mediated stripping of eIF4G:E from uncapped locations on the mRNA. Error bars represent the standard deviations of replicate measurements, which are shown as points. The baselines (eIF4A_WT_^1^ and photobleaching) are plotted from our previously published work^24^.

**Extended Data Figure 2.**
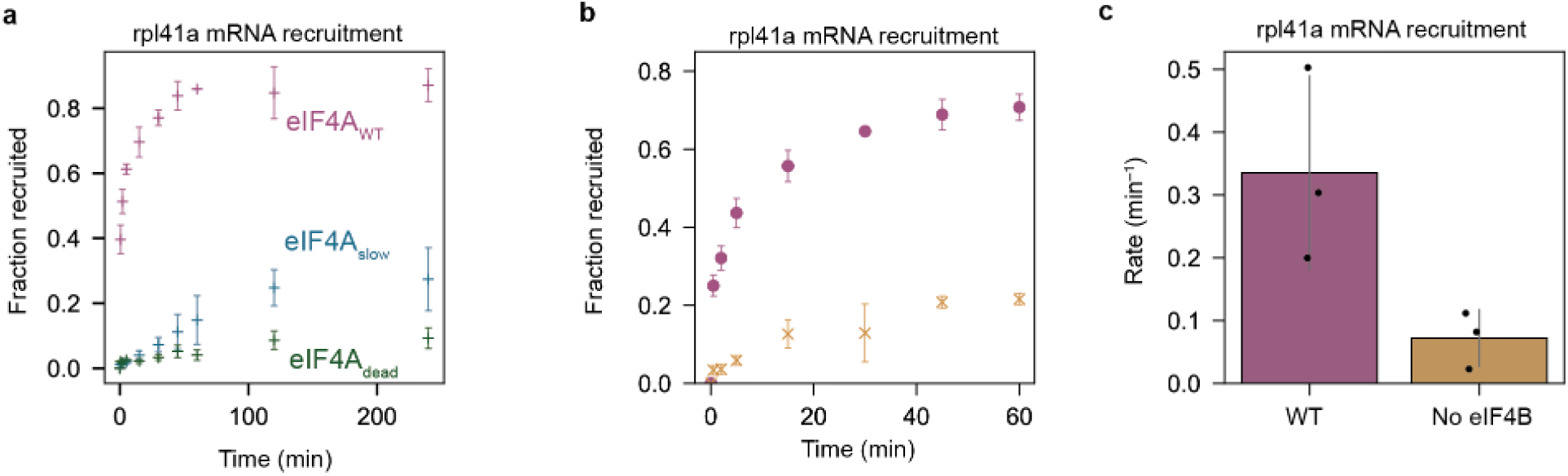
ATP hydrolysis by eIF4A and eIF4B are required for efficient cap-dependent mRNA recruitment. **(a)** Ensemble mRNA recruitment assays monitoring recruitment of capped rpl41a mRNA in the presence of wild-type eIF4A, eIF4A_slow_, or eIF4A_dead_. Compromising ATP hydrolysis progressively impairs cap-dependent mRNA recruitment. Points represent the mean, and error bars represent the standard deviations of replicate measurements. **(b)** Ensemble mRNA recruitment assays monitoring recruitment of capped rpl41a mRNA in the presence or absence of eIF4B. Omission of eIF4B substantially slows mRNA recruitment. Points represent the mean, and error bars represent the standard deviations of replicate measurements. **(c)** Quantification of recruitment rates measured in (b). Error bars represent the standard deviations of replicate measurements, which are shown as points.

**Extended Data Figure 3.**
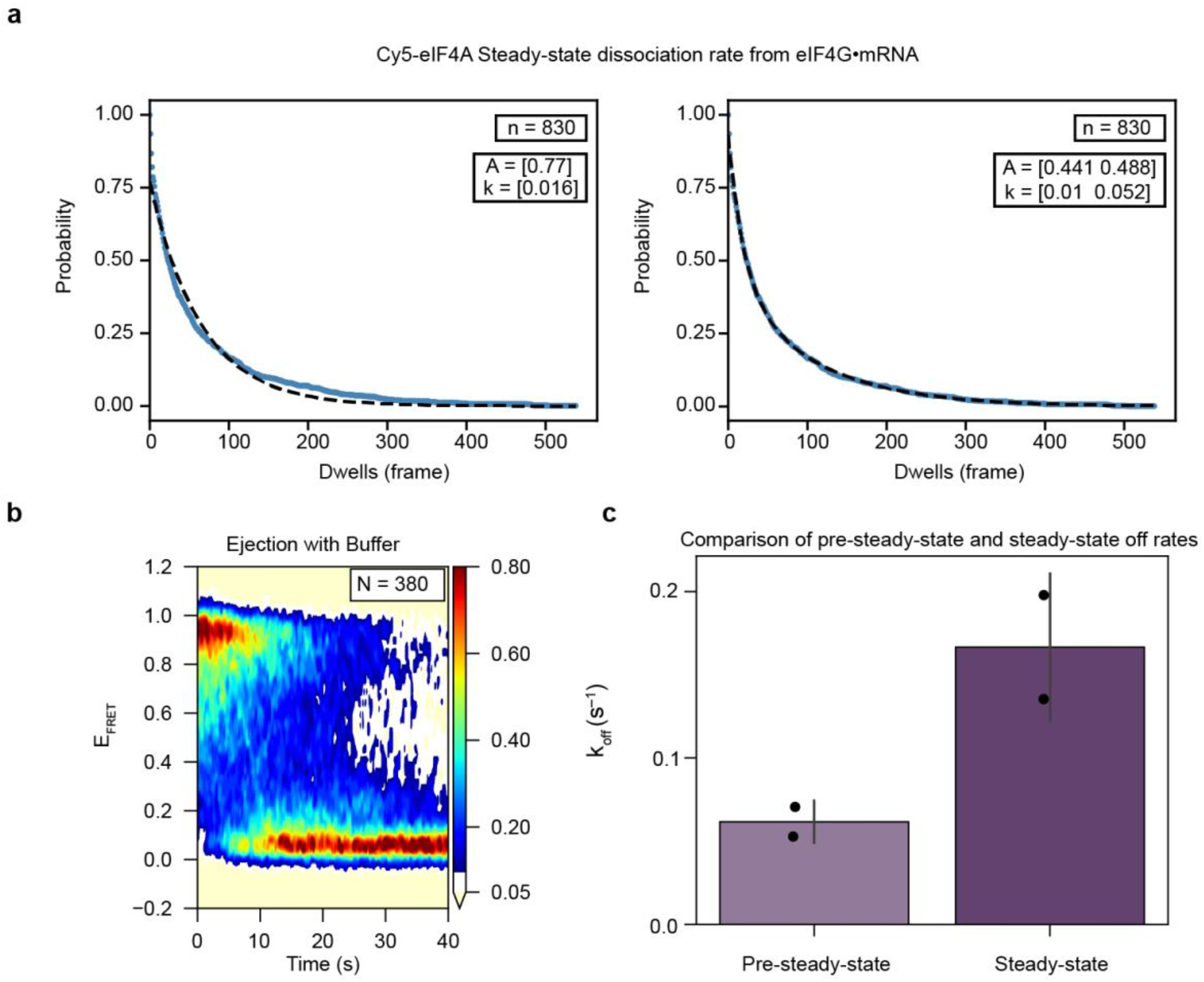
eIF4A facilitates its own dissociation from eIF4G:E•mRNA complexes. **(a)** Survival analysis of steady-state Cy5-eIF4A dissociation events from eIF4G:E•mRNA complexes. The dwell-time distribution is better described by a bi-exponential model than by a single-exponential model, indicating multiple dissociation pathways during steady-state turnover. **(b)** Surface contour plot showing dissociation of pre-bound Cy5-eIF4A from eIF4G:E•mRNA complexes following removal of unbound protein by buffer exchange. **(c)** Comparison of pre-steady-state and steady-state dissociation rates. Pre-steady-state dissociation is substantially slower than the population-weighted steady-state dissociation rate, consistent with eIF4A facilitating its own dissociation through rebinding events under steady-state conditions. Error bars represent the standard deviations of replicate measurements, which are shown as points.

**Extended Data Figure 4.**
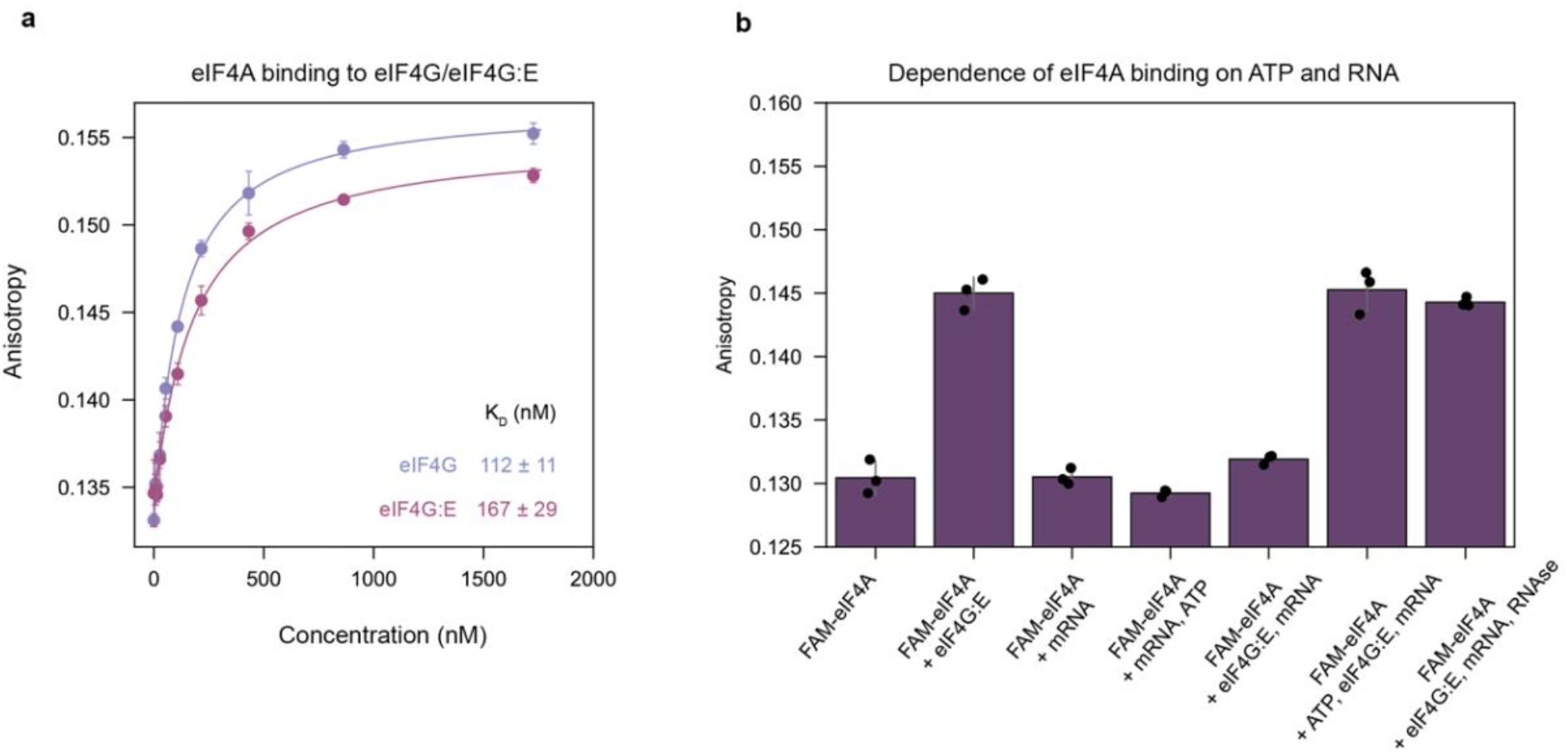
ATP-dependent assembly of eIF4F on mRNA measured by fluorescence anisotropy. **(a)** Fluorescence anisotropy titrations measuring binding of FAM-eIF4A to eIF4G or eIF4G:E in the absence of mRNA. Both eIF4G and eIF4G:E bind eIF4A with high affinity under these conditions. Curves represent fits to a single-site binding model, and the corresponding dissociation constants (*K*_D_) are indicated. Points represent the mean, and error bars represent the standard deviations of replicate measurements. **(b)** Endpoint fluorescence anisotropy measurements examining assembly of eIF4A with eIF4G:E under different nucleotide and RNA conditions. In the absence of mRNA, eIF4G:E increases the anisotropy of FAM-eIF4A, consistent with complex formation. Addition of mRNA abolishes this increase in the absence of ATP, whereas anisotropy is restored when both mRNA and ATP are present. Similarly, treatment with benzonase restores the increase in anisotropy in the absence of ATP. These results independently validate that productive assembly of the ATP-bound eIF4F•mRNA complex requires ATP. Error bars represent the standard deviations of replicate measurements, which are shown as points.

**Extended Data Figure 5.**
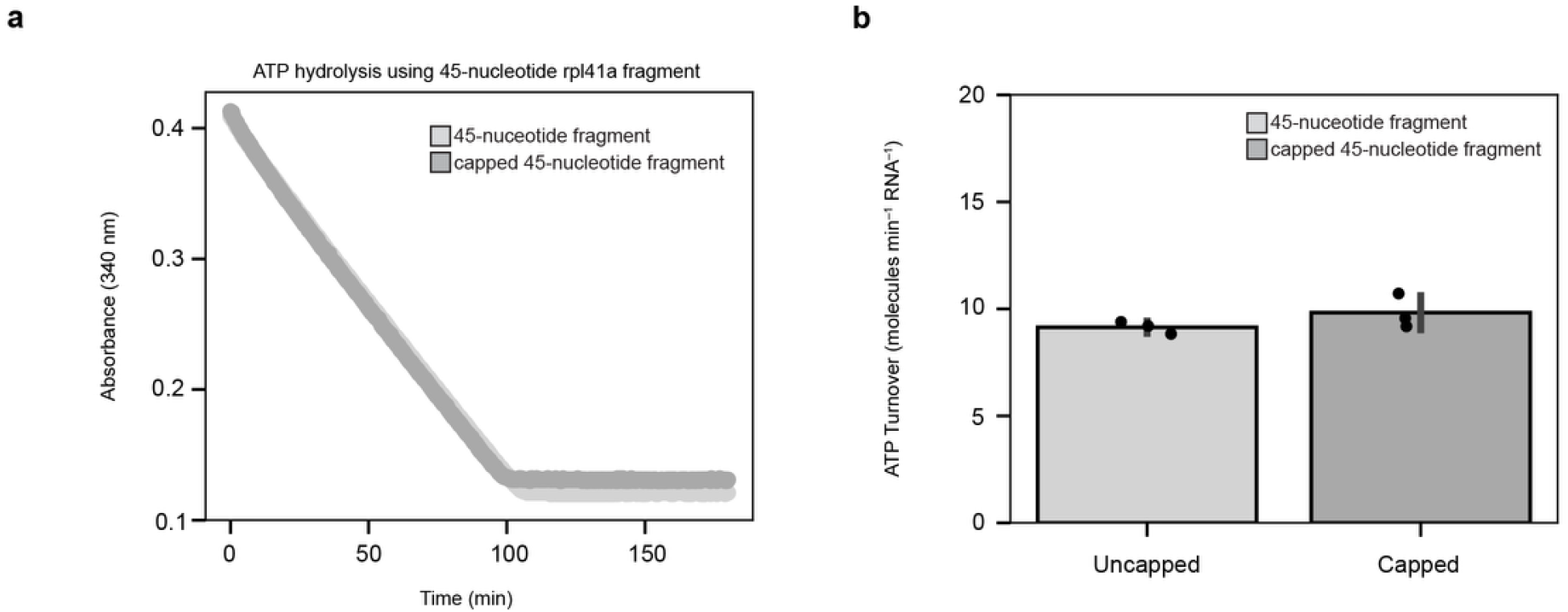
Cap recognition does not alter the ATP hydrolysis rate of eIF4F. **(a)** Representative NADH-coupled ATPase assays comparing ATP hydrolysis by eIF4F assembled on uncapped and capped versions of a 45-nucleotide rpl41a mRNA fragment. The short mRNA construct was designed to accommodate only a single eIF4F complex and to ensure cap-proximal binding when capped. ATP hydrolysis was monitored through loss of NADH absorbance at 340 nm. **(b)** Quantification of ATP hydrolysis rates measured in (a). ATP hydrolysis by eIF4F was indistinguishable on capped and uncapped mRNA substrates, demonstrating that cap recognition by eIF4E does not modulate the ATP hydrolysis activity of eIF4F. Error bars represent the standard deviations of replicate measurements, which are shown as points.

**Extended Data Figure 6.**
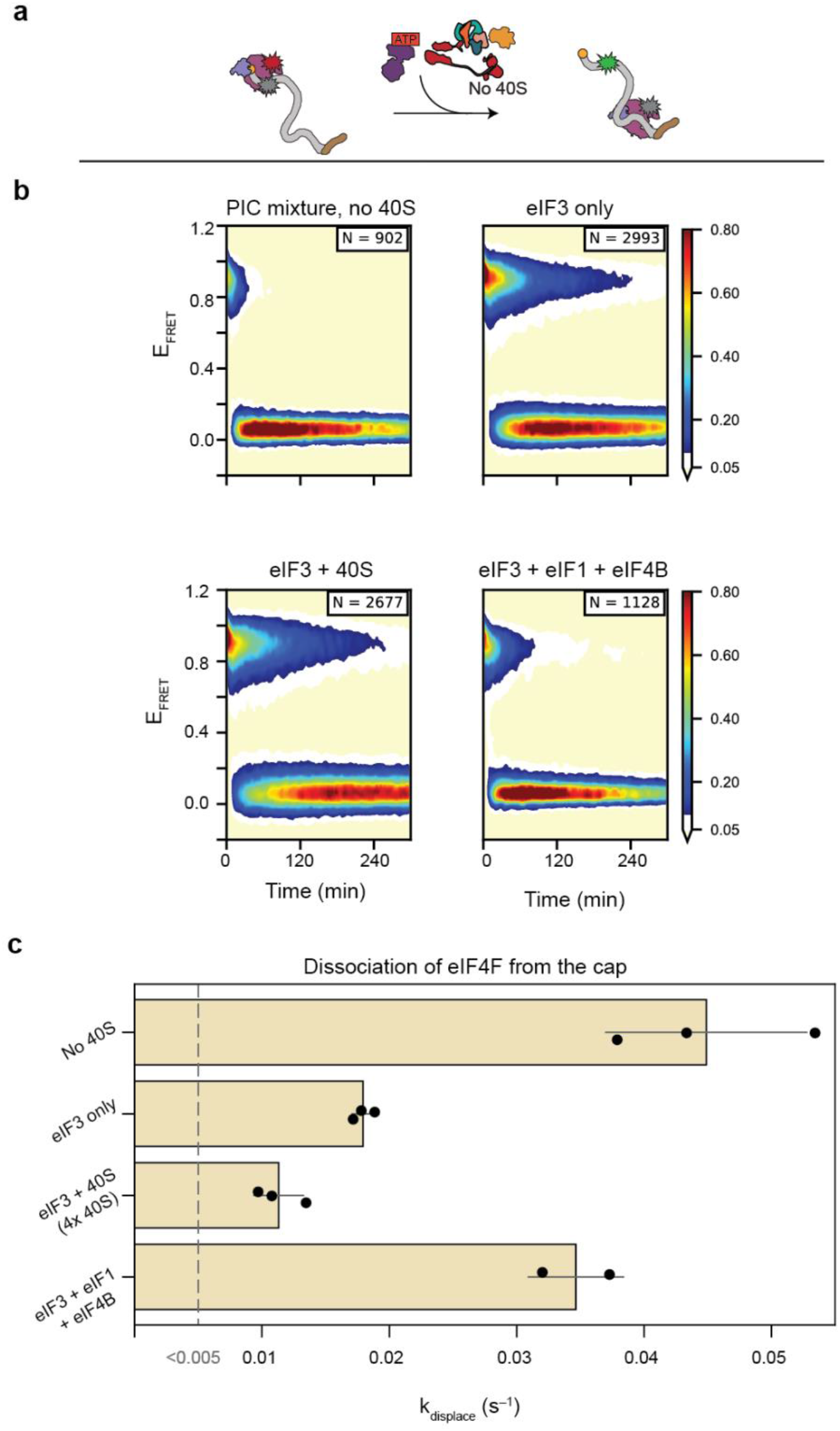
eIF3 stimulates eIF4F recycling independently of the 40S subunit, whereas the 40S subunit suppresses eIF3-dependent recycling in the absence of eIF4B. **(a)** Schematic of the single-molecule assay used to measure ATP-hydrolysis-dependent recycling of cap-trapped Cy5-eIF4G:E. Cy5-eIF4G:E was first allowed to associate with capped rpl41a mRNA before injection of solutions containing eIF4A, ATP•Mg, and the indicated initiation factors. **(b)** Surface contour plots showing dissociation of cap-associated Cy5-eIF4G:E under the indicated conditions. Omission of the 40S subunit from the PIC mixture does not impair eIF4F recycling, whereas eIF3 alone stimulates recycling less efficiently than the complete PIC mixture. Addition of excess 40S subunits suppresses eIF3-dependent recycling, and efficient recycling is restored by inclusion of eIF4B. **(c)** Quantification of eIF4G:E dissociation rates measured in (b). These results demonstrate that eIF3 can stimulate eIF4F recycling independently of the 40S subunit, but that 40S-bound eIF3 requires eIF4B to efficiently promote ATP-hydrolysis-dependent recycling of eIF4F. Error bars represent the standard deviations of replicate measurements, which are shown as points.

**Extended Data Figure 7.**
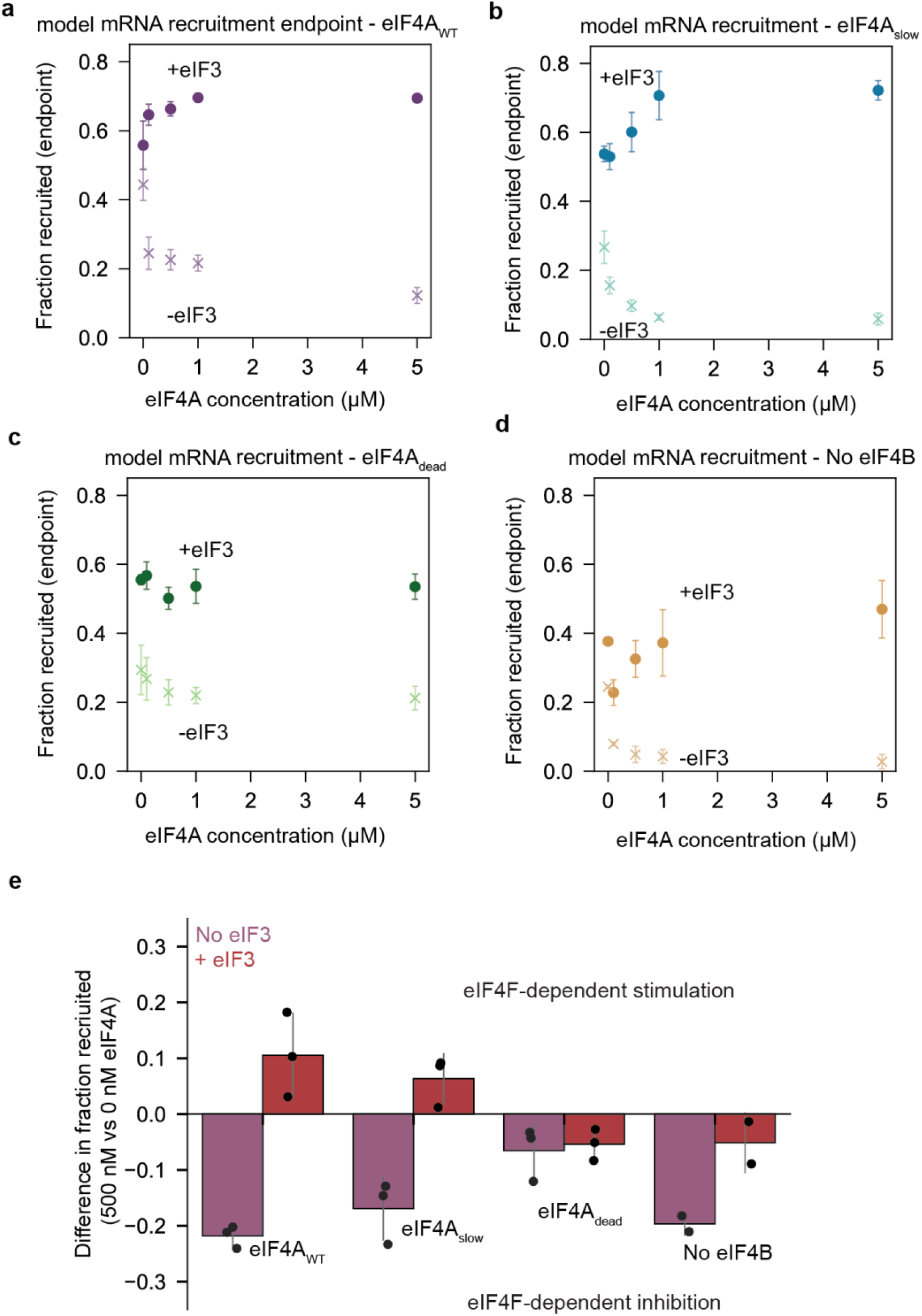
eIF3 and eIF4B convert cap-trapped eIF4F from an inhibitor into a stimulator of model mRNA recruitment. Endpoint mRNA recruitment assays using a capped poly(CAA) model mRNA. Recruitment was measured as a function of eIF4A concentration in the presence (+eIF3) or absence (−eIF3) of eIF3 using **(a)** wild-type eIF4A, **(b)** eIF4A_slow_, **(c)** eIF4A_dead_, or **(d)** wild-type eIF4A in reactions lacking eIF4B. Because efficient recruitment of eIF4G:E to the cap requires eIF4A in this system, increasing eIF4A concentration promotes formation of cap-trapped eIF4F complexes. In the absence of eIF3, cap-trapped eIF4F inhibits mRNA recruitment, consistent with an inability to recycle from the cap. In the presence of eIF3, however, this inhibition is converted into stimulation, indicating that eIF3 promotes productive eIF4F recycling during mRNA recruitment. This stimulation is reduced when ATP hydrolysis is compromised by eIF4A_slow_ or eIF4A_dead_ and is abolished in the absence of eIF4B. Points represent the mean, and error bars represent the standard deviations of replicate measurements. **(e)** Quantification of the effect of cap-trapped eIF4F on mRNA recruitment, calculated as the difference in endpoint recruitment between reactions containing 500 nM eIF4A and reactions lacking eIF4A. Negative values indicate eIF4F-dependent inhibition of recruitment, whereas positive values indicate eIF4F-dependent stimulation. Together, these results demonstrate functional cooperativity between eIF3, eIF4B, and ATP-hydrolysis-dependent recycling of eIF4F during mRNA recruitment. Error bars represent the standard deviations of replicate measurements, which are shown as points.

**Extended Data Table 1.** Summary of eIF4G:E cap binding lifetimes.

| Summary of eIF4G:E cap binding lifetimes |  |  |  |  |
| --- | --- | --- | --- | --- |
| Injection contains | Lifetime (s) | $k_{\text{displace}}$ ($\text{s}^{-1}$) | Relative instability | $K_D$ (pM)<sup>a</sup> |
| **ATPase Activity<sup>b</sup>** |  |  |  |  |
| Buffer only | 181 ± 19 | 0.0055 ± 0.0006 | 1.0 | 17 ± 2 |
| eIF4A | 114 ± 22 | 0.0088 ± 0.0017 | 1.6 | 28 ± 5 |
| eIF4A + PIC<sup>b</sup> | 24 ± 5 | 0.0411 ± 0.0084 | 7.4 | 130 ± 27 |
| eIF4A + PIC no ATP<sup>b</sup> | 188 ± 11 | 0.0053 ± 0.0003 | 1.0 | 17 ± 1 |
| eIF4A<sub>slow</sub> + PIC<sup>b</sup> | 168 ± 2 | 0.006 ± 0.0001 | 1.1 | 19 ± 0 |
| eIF4A<sub>dead</sub> + PIC<sup>b</sup> | 183 ± 17 | 0.0055 ± 0.0005 | 1.0 | 17 ± 2 |
| **Effect of eIF3<sup>c</sup>** |  |  |  |  |
| eIF4A + PIC no eIF3<sup>c</sup> | 101 ± 11 | 0.0099 ± 0.0011 | 1.8 | 31 ± 3 |
| eIF4A + eIF3 | 56 ± 3 | 0.0179 ± 0.0009 | 3.2 | 57 ± 3 |
| eIF4A + eIF3 + 4B + 1 | 29 ± 3 | 0.0347 ± 0.0037 | 6.3 | 110 ± 12 |
| eIF4A + eIF3 + 40S<sup>c</sup> | 88 ± 15 | 0.0113 ± 0.0019 | 2.0 | 36 ± 6 |
| eIF4A + PIC<sup>c</sup> | 29 ± 0 | 0.0343 ± 0.0004 | 6.2 | 109 ± 2 |
| eIF4A + eIF3 + 4B + 40S<sup>c</sup> | 36 ± 2 | 0.0275 ± 0.0018 | 5.0 | 87 ± 6 |
| **PIC eIF Dropouts<sup>c</sup>** |  |  |  |  |
| No 40S | 22 ± 4 | 0.0449 ± 0.0079 | 8.1 | 142 ± 25 |
| No TC<sup>c</sup> | 28 ± 2 | 0.0352 ± 0.0027 | 6.4 | 111 ± 9 |
| No 4B<sup>c</sup> | 74 ± 3 | 0.0136 ± 0.0006 | 2.5 | 43 ± 2 |
| No TC, 4B<sup>c</sup> | 92 ± 18 | 0.0109 ± 0.0021 | 2.0 | 35 ± 7 |

<sup>a</sup>Affinities calculated using a previously determined $k_{\text{on}}$ of $316 \pm 5$
<sup>b</sup>Concentration of 40S subunits was 100 nM, which is half of the 200 nM eIF3 used
<sup>c</sup>Concentration of 40S subunits was 400 nM, which is twice the 200 nM eIF3 used

